# Evolutionary compromises to environmental toxins: ammonia and urea tolerance in *Drosophila suzukii* and *Drosophila melanogaster*

**DOI:** 10.1101/124685

**Authors:** Virginia Belloni, Alessia Galeazzi, Giulia Bernini, Mauro Mandrioli, Elisabetta Versace, Albrecht Haase

## Abstract

The invasive species *Drosophila suzukii* has evolved morphological and behavioral adaptations to lay eggs under the skin of fresh fruits. This results in severe damage of a wide range of small and stone fruits, thus making this species a serious agricultural and economical threat.

*Drosophila suzukii* females typically lay few eggs per fruit, preferring not infested fruits. Hence larvae are exposed to a reduced amount of nitrogenous waste products. On the contrary, the innocuous *Drosophila melanogaster* lays eggs on fermented fruits already infested by conspecifics, with larvae developing in a crowded environment characterized by accumulation of nitrogenous waste such as ammonia and urea. Given these differences in oviposition and larval ecological niche, we expected different behavioral and physiological mechanisms in the two species to cope with nitrogenous waste. We investigated the impact of different concentrations of ammonia and urea on fecundity and larval development in both species. Females and larvae of *D. suzukii* showed a greater sensitivity to high concentration of both compounds, with a dramatic decrease in fecundity and egg viability.

To better understand the pathways underlying these differences, we evaluated the effect on ornithine aminotransferase and glutathione-S-transferase, two enzymes involved in nitrogen metabolism and stress response that are expressed during larval development. Under ammonia and urea exposure, the expression of these enzymes was significantly reduced in *D. suzukii*.

The fact that *D. suzukii*’s shift from rotten to fresh fruit as oviposition and larval substrate resulted in less efficient detoxifying and excretory mechanisms represents a potential approach for its control. Fecundity and larval development are in fact dramatically impaired by nitrogen waste products. These findings can help in planning effective strategies of sustainable pest management that targets both females and larvae.

## Introduction

In the last decade, growing concern has turned on the invasive pest *Drosophila suzukii*, a serious agricultural and economical threat (Bolda, Goodhue & Zalom 2010; De Ros *et al.* 2012). This species is native to Asia and has recently invaded western countries, with rapidly expanding range in America and Europe (Rota-Stabelli, Blaxter & Anfora 2013; Walsh *et al.* 2011). Differently from other *Drosophila* species, that attack overripe and decaying fruits, females of *D. suzukii* lay eggs under the skin of fresh fruits, through a serrated ovipositor. Therefore, larval development and exposure to pathogens result in damage of a wide range of small and stone fruits (Goodhue *et al.* 2011; Rota-Stabelli, Blaxter & Anfora 2013). To date, most research on *D. suzukii* has focused on adults (Crava *et al.* 2016; Hamby & Becher 2016; Gong *et al.* 2016; Keesey, Knaden & Hansson 2015; Rossi-Stacconi *et al.* 2016), with little attention to larvae. A more comprehensive understanding of the ecology and biology of this species is important to develop management strategies and to successfully minimize its spread and impact (Cini 2012; Dreves 2011).

The innocuous *Drosophila melanogaster* lays eggs in rotten fruits and larvae develop in a crowded environment, rich in bacteria, mold, and yeast (Becher *et al.* 2012; Zhu, Park & Baker 2003). Females of this species have a gregarious tendency in selecting the oviposition site and prefer to lay eggs where other larvae are present (Sarin & Dukas 2009; del Solar & Palomino 1966). High larval density combined with microorganism metabolic activity and protein-rich microbial community (Begon 1982; Chandler, Eisen & Kopp 2012) result in accumulation of nitrogen waste products such as ammonia and, at lower extent, urea (Botella *et al.* 1985; Mueller 1995), that are relatively toxic when concentrated in organism tissues (David *et al.* 1999; Maas, Seibel & Walsh 2012). In *D. melanogaster*, high concentrations of dietary urea and ammonia have been associated with a decrease in female fecundity (e.g. Joshi *et al.* 1997, Min *et al.* 2013), a decline in egg-to-adult viability, as well as an increase in developmental time (Borash *et al.* 2000; Shiotsugu *et al.* 1997). Due to the limited mobility of larvae (Durisko *et al.* 2014; Philippe *et al.* 2016), behavioral avoidance cannot prevent larval exposure to environmental toxins accumulating in the food, but physiological mechanisms help larvae to cope with toxic compounds (Heinstra *et al.* 1989; Wilson 2001). *D. melanogaster* populations reared under crowded larval conditions develop greater competitive ability (Borash *et al.* 2000; Sarangi *et al.* 2016) and increased resistance to both urea and ammonia (Borash *et al.* 2000; Etienne, Fortunat & Pierce 2001). While the response of *D. melanogaster* to high levels of urea and ammonia has been studied (Borash *et al.* 2000; Etienne, Fortunat & Pierce 2001), little is known on the effects in *D. suzukii*. This species occupies a unique ecological niche compared to other drosophilids, since larval development occurs in fresh fruits (Rota-Stabelli, Blaxter & Anfora 2013), rich in water (Ishida, Koizumi & Kano 1994) and relatively poor in microorganisms, due to the skin barrier (Tournas & Katsoudas 2005). Moreover, females of *D. suzukii* tend to lay few eggs per fruit (Burrack *et al.* 2013; Yu, Zalom & Hamby 2013), resulting in a moderate larval density and, as a consequence, a low level of waste products. Thus, we hypothesised to observe differences in the behavioral and physiological responses of *D. melanogaster* and *D. suzukii* to nitrogenous waste products, as an effect of adaptation to different ecological niches

To evaluate tolerance capacity for nitrogenous compounds in *D. suzukii* compared to *D. melanogaster*, we investigated the effect of different concentrations of urea and ammonia on female fecundity, under no-choice and choice conditions, and on larval development in both species. We further studied potential differences in the expression of ornithine aminotransferase and glutathione-S-transferase, enzymes involved in metabolic and detoxifying pathways. Ornithine aminotransferase is an enzyme highly expressed in third instar larvae of *D. melanogaster*, playing a crucial role in amino acids metabolism and nitrogen homeostasis (Ventura *et al.* 2008; Yoshida, Juni & Hori 1997). Glutathione-S-transferase is an enzyme involved in insect resistance to endogenous and xenobiotic compounds and in protection against oxidative stress (Hamby *et al.* 2013; Perry, Batterham & Daborn 2011), which is expressed in the larval midgut of *D. melanogaster* (Li *et al.* 2008).

## Materials and methods

### Insect strains and rearing

We used adult flies of *Drosophila melanogaster* from 50 lines of the DGRP population (Mackay *et al.* 2012), a collection of inbred isofemale lines originally collected in Raleigh, US (Ayroles *et al.* 2009). Lines were obtained from the Bloomington Drosophila Stock Center (Indiana University, Bloomington, US) and represent a spectrum of natural occurring genetic variation. The same isofemale lines were tested in all treatment groups. The *Drosophila suzukii* flies used in this study were originally collected in the Trentino area, Italy, and maintained under the same laboratory conditions of the *Drosophila melanogaster* population for several generations. All flies were raised on a standard Drosophila diet (see Appendix 1. in Supporting Information) at 25 ± 1 °C with 65 ± 1% relative humidity with a light:dark cycle of 14:10 h.

### Chemicals

Ammonium chloride (NH_4_Cl; purity≥99.5%) was purchased from Carl Roth (Karlsruhe, Germany); urea (ACS reagent 99-100.5%) was purchased from Sigma-Aldrich (St. Louis, USA). Propionic acid was obtained from Carlo Erba Reagents (Milan, Italy) and methyl 4-hydroxybenzoate (99%) was purchased from Acros Organics (Morris Plains, US).

Ammonium chloride and urea (pH ~5.5) were added to the standard food medium, after it had cooled down to 48 °C, and antimicrobial agents were added to the food. In order to homogenize the mixture, the supplemented media was placed upon a magnetic stirring apparatus, which rapidly stirred the mixture as it was dispensed into polypropylene vials (25 x 95 mm) or Ø 90 mm petri dishes.

### Female fecundity and larval development in a no-choice assay

Newly eclosed flies were transferred to fresh food vials, and were maintained under standard conditions until tested. Females (5-6 days old) of about the same size were individually assigned, using light CO_2_ anesthesia, to a vial with 5 ml of standard diet with one of the following supplements: urea at 25 mM (UL=urea low concentration) or 250 mM (UH=urea high concentration) or ammonium chloride at 25 mM (AL=ammonia low concentration) or 250 mM (AH=ammonia high concentration), or no supplements added to the food (CTRL). A pinch of active yeast was sprinkled on the food to stimulate oviposition. After 24 h, females were removed from the vials and eggs counted under an optical microscope. In this assay we quantified fecundity as the number of eggs laid within 24 hours. One day later, presence of larvae and their conditions (alive/dead, 1^st^/2^nd^ instar) were recorded. After the first pupa appeared, pupae were counted and number and times of pupation were recorded twice per day, at 10 a.m. and 5 p.m., for the following five days. As development was expected to be slower on both ammonia- and urea-supplemented foods, the scores continued during the following days. The larval developmental time has been considered as the number of hours occurring from hatching to pupation. From the beginning of adult emergence, flies were collected, using CO_2_ anesthesia, and the number of adults and their sex were recorded for each vial. Data were collected every morning (10 a.m.) and checks ceased when in a period of 48 h no more flies had emerged from a given population. The experiment was replicated 6 times for a total sample size of 32 females for each treatment group. Larval development (time to pupation and number of pupae) and viability (eggs-to-pupae and egg-to-adults) were evaluated on 25 vials for each treatment group.

### Female fecundity in a choice assay

Oviposition behavior was tested both under no-choice and dual choice conditions to control for interaction between environmental cues and treatment (Lihoreau et al. 2016; Sheeba, Madhyastha & Joshi 1998). Flies were tested in a cage where both a control and an experimental medium were provided. The oviposition substrates consisted of Ø 90 mm petri dishes filled with 20 ml of standard food (CTRL), or with standard food and one of the following supplements: urea at 25 mM (UL) or 250 mM (UH), or ammonium chloride at 25 mM (AL) or 250 mM (AH). Sprinkles of active yeast were added to the food to stimulate oviposition.

Freshly eclosed flies were transferred to fresh food vials and were maintained at standard conditions until tested. Ten females (5-6 days old) of each species were collected, using CO_2_ anesthesia, and transferred to a bug dorm insect-rearing cage (30x30x30 cm; BugDorm-1, MegaView Science Taichung, Taiwan) with the control petri dish in one corner, and the petri dish with supplemented food on the opposite side. After 24 hours, the petri dishes were collected and the eggs counted under an optical microscope. In this assay we quantified fecundity as the number of eggs laid in 24 hours. The experiment included 10 replicates for each condition.

### Semiquantitative analysis of the expression of genes coding for metabolic and detoxifying enzymes

Expression of ornithine aminotransferase (OAT) and glutathione-S-transferase D2 (gstD2) and D4 (gstD4) was assayed by RT-PCR in third instar larvae that had developed in standard food (CTRL) or in standard food and one of the following supplements: urea at 25 mM (UL) or 250 mM (UH) or ammonium chloride at 25 mM (AL) or 250 mM (AH).

A total sample size of about 15 larvae per treatment group was collected. We had no samples from the AH group, because no larvae arrived at the third instar stage. Total RNA extracted from samples using a TRIreagent:chloroform (Sigma-Aldrich, St. Louis, US) extraction, performed according to the manufacturer’s instructions. RNA samples were quantified using a Nanodrop spectrophotometer (ThermoFisher Scientific, Waltham, US) and reverse-transcribed to cDNA using the Revert Aid First Strand cDNA Synthesis Kit (ThermoFisher Scientific, Waltham, US) with specific primers (table S1.). PCR products were analysed by gel electrophoresis in a 1% ethidium bromide-stained agarose gel. Gel documentation was collected using a “Gel Doc XR”, digitally evaluated with “Quantity One” (Bio-Rad Lab., Milano, Italy) and normalized to the correspondent signals for tubulin.

### Statistical analysis

For each species, non-parametric data related to eggs number, eggs-to-pupae and eggs-to-adults viability, sex-ratio, and densitometry for enzymes expression were tested by a Kruskal-Wallis test. To compare the effect between species, the number of eggs was normalized relative to the control, and was analyzed with a Mann-Whitney test. Data related to larval developmental time (from hatching to pupation) were averaged for each vial and tested by a one-way analysis of covariance (ANCOVA) with Treatment as factor and number of eggs as covariate. A Bonferroni correction was applied when post hoc multiple comparisons were performed.

The number of eggs laid in the preference test was analyzed with a Wilcoxon signed-rank test to compare conditions (CTRL versus Treatment). A *p*-value of less than 0.05 was considered significant. All statistical analyses were carried out using SPSS version 17 (IBM, Armonk, US).

## Results

### Female fecundity and larval development in no-choice assay

No significant difference in female fecundity was observed at low concentration of ammonia and urea with respect to the control. At high concentration, ammonia (AH) reduced significantly the number of eggs in *D. melanogaster* (*χ*^2^(4)=22.89, *p*<0.001, see Fig. 1a). However, high concentrations of both urea (UH) and ammonia (AH) decreased female fecundity in *D. suzukii* (*χ*^2^(4)=70.77, *p*<0.001, see Fig. 1b). When comparing the effects between species, *D. suzukii* females showed a greater sensitivity to the treatment (AH: *U*=195, *p*<0.001; UH: *U*=237, *p*<0.001, see Appendix 1).

Twenty-four hours after the start of hatching, first and second instar larvae were present in all treatment groups except for AH of *D. suzukii* (Fig. 2a, b). In this condition, larvae died soon after hatching, and many of them were observed on the wall of the vial, suggesting a possible escaping behavior. In *D. melanogaster*, larval development was affected by exposure to high concentration of urea (140±3 hours) and ammonia (130±2 hours) with respect to the control (106±2 hours), with a significant delay in pupation (Treatment: *F*(4,119)=68.24, *p*<0.001). These durations did not depend on the number of eggs (Treatment*Number of eggs: *F*(4,119)=1.54, *p*=0.19, see Appendix 2). However, when considering eggs-to-pupae viability, a significant decrease was observed only in UH (*χ*^2^(4)=25.76, *p*=0.01, see Fig. 3a), while no difference was found under ammonia exposure. In *D. suzukii*, the effect of the treatment was even stronger, with no pupae in the AH group, and only a few pupae under high concentration of urea (*χ*^2^(4)=78.16, *p*<0.001, see Fig 3b), accompanied by a visible delay.

**Figure 1.**
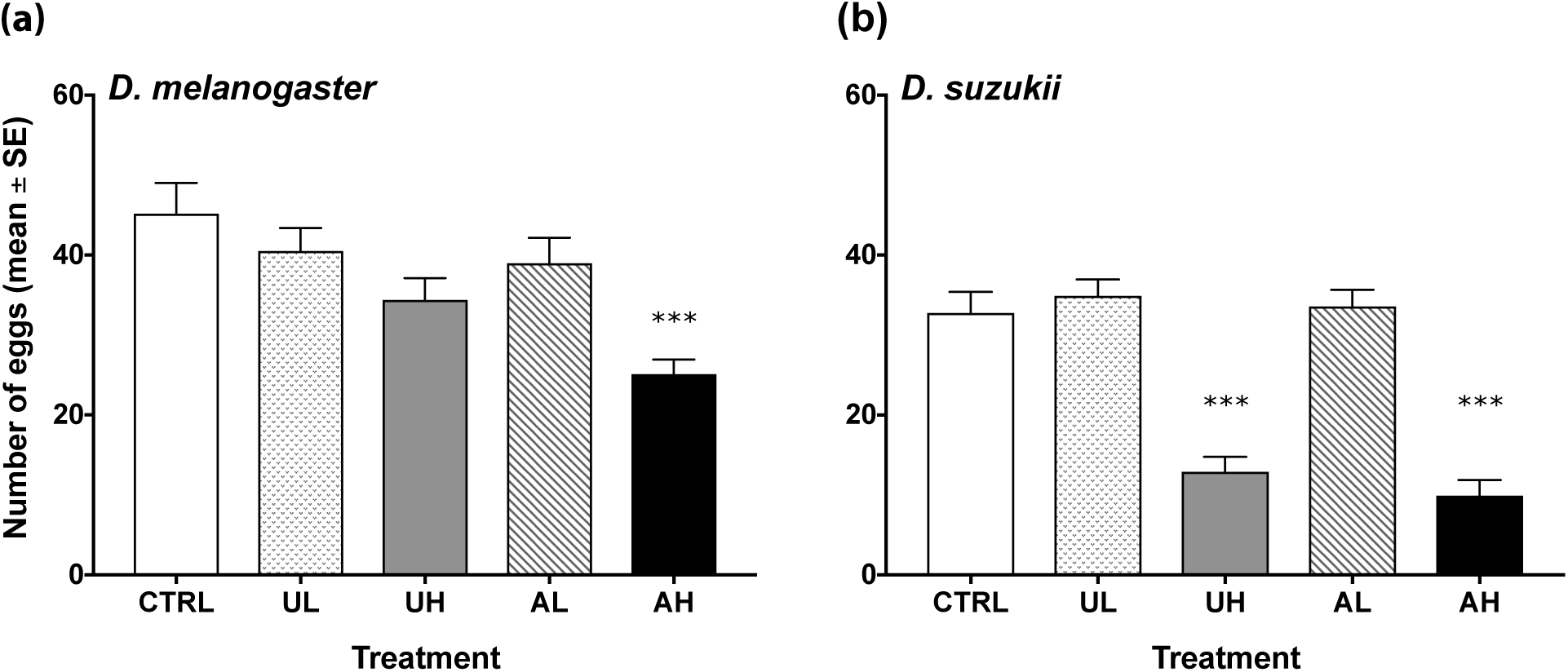
Eggs laid during a 24-hour period by females exposed to standard food and standard food supplemented with urea and ammonia. CTRL: standard food; UL: standard food with 25 mM of urea; UH: standard food with 250 mM of urea; AL: standard food with 25 mM of ammonium chloride; AH: standard food medium with 250 mM of ammonium chloride. Mean ± standard error (SE) are shown, ***=*p*<0.001.

**Figure 2.**
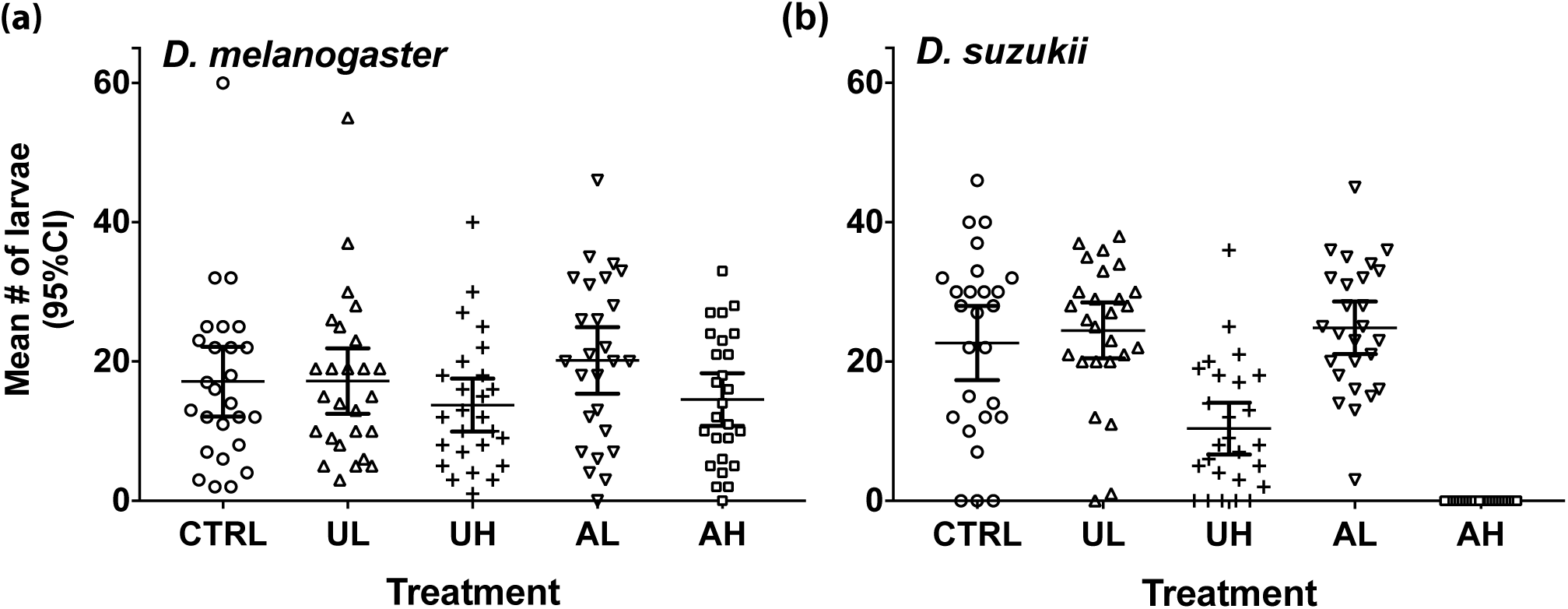
Estimated number of alive larvae 24 hours after hatching in standard food and in standard food supplemented with urea or ammonia. CTRL: standard food; UL: standard food with 25 mM of urea; UH: standard food with 250 mM of urea; AL: standard food with 25 mM of ammonium chloride; AH: standard food medium with 250 mM of ammonium chloride. Horizontal bars indicate mean and 95% confidence interval.

**Figure 3.**
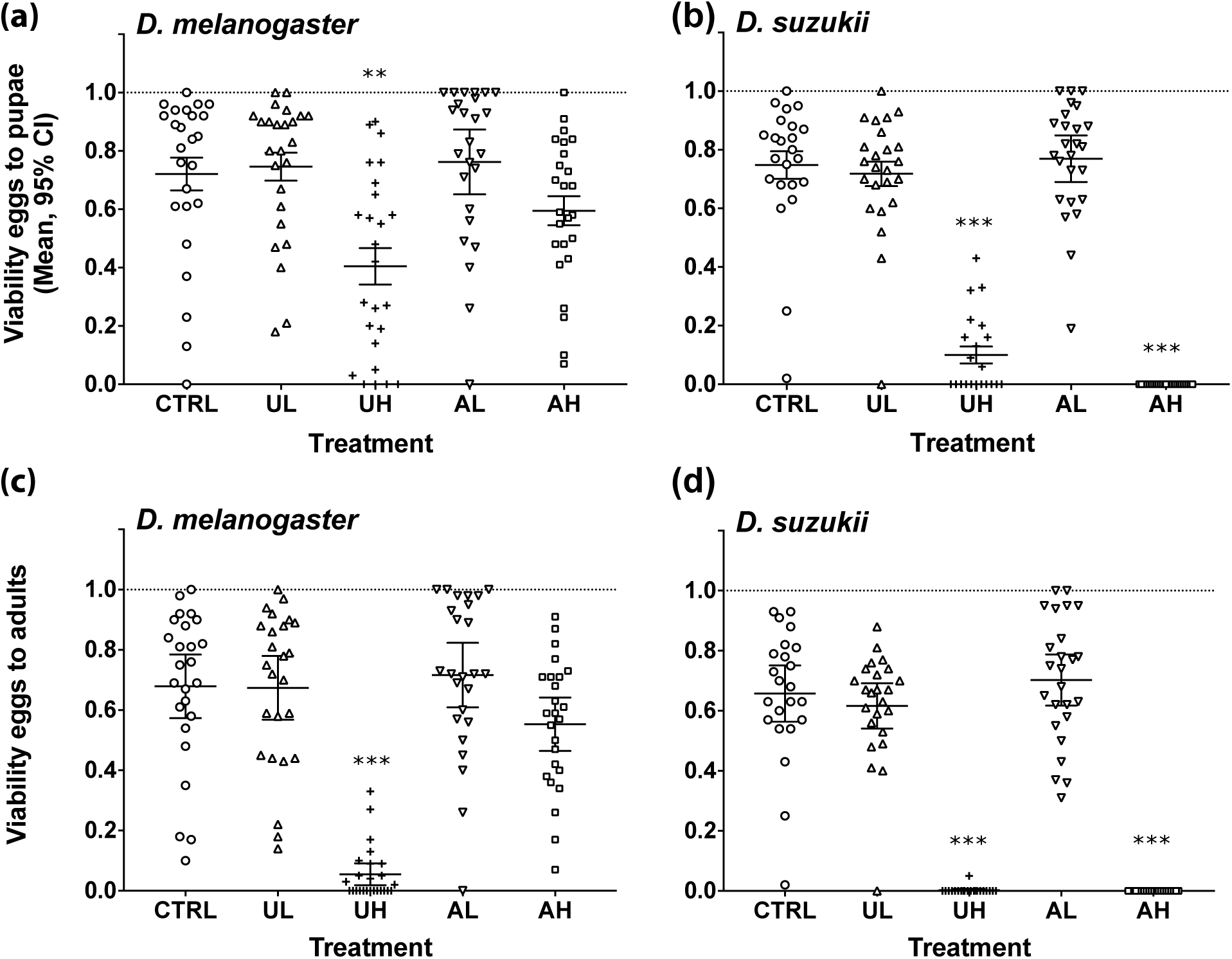
Viability eggs-to-pupae (a, b) and eggs-to-adults (c, d) in standard food and standard food supplemented with dietary urea and ammonia. CTRL: standard food; UL: standard food with 25 mM of urea; UH: standard food with 250 mM of urea; AL: standard food with 25 mM of ammonium chloride; AH: standard food medium with 250 mM of ammonium chloride. Horizontal bars indicate mean and 95% confidence interval. **=*p*≤0.01, ***=*p*<0.001.

Finally, pupariation and emergence of adult flies was strongly impaired by a high concentration of urea in both species (*D. melanogaster*: *χ*^2^(4)=60.19, *p*<0.001; *D. suzukii: χ*^2^(3)=48.94, *p*<0.001, see Fig. 3c, d), while eggs-to-adults viability was not significantly affected by high concentration of ammonia in *D. melanogaster* (see Fig. 3c). No difference in adult sex ratio was observed among groups (*D. melanogaster*: *χ*^2^(4)=1.69, *p*=0.8; *D. suzukii*: *χ*^2^(2)=1.72 *p*=0.4).

### Female fecundity in a choice assay

Oviposition preference was tested in a choice assay between experimental substrates (supplemented with urea or ammonia) and control substrates. Females did not show significant egg laying preferences between control food and food supplemented with low concentration of urea (*D. melanogaster* Z(9)=-0.76, *p*=0.5, *D. suzukii* Z(9)=-1.78, *p*=0.08) and ammonia (*D. melanogaster* Z(9)=-1.07, *p*=0.3, *D. suzukii* Z(9)=-1.63, *p*=0.1, see Fig. 4a,b) When the concentration of ammonia was increased, females displayed a strong aversion and only about 11% of the eggs where laid in the AH substrate in *D. melanogaster* (Z(9)=-2.70, *p*<0.01, see Fig. 4a) and 9% in *D. suzukii* (Z(9)=-2.80, *p*<0.01, see Fig. 4b). Interestingly, oviposition preference was unaffected by high concentration of urea in *D. melanogaster* (53% of the eggs where laid in the UH substrate), whereas *D. suzukii* females laid significantly less in the UH site (29%) than in the urea-free medium (Z(9)=-2.09, *p*<0.05, see Fig. 4a,b).

**Figure 4.**
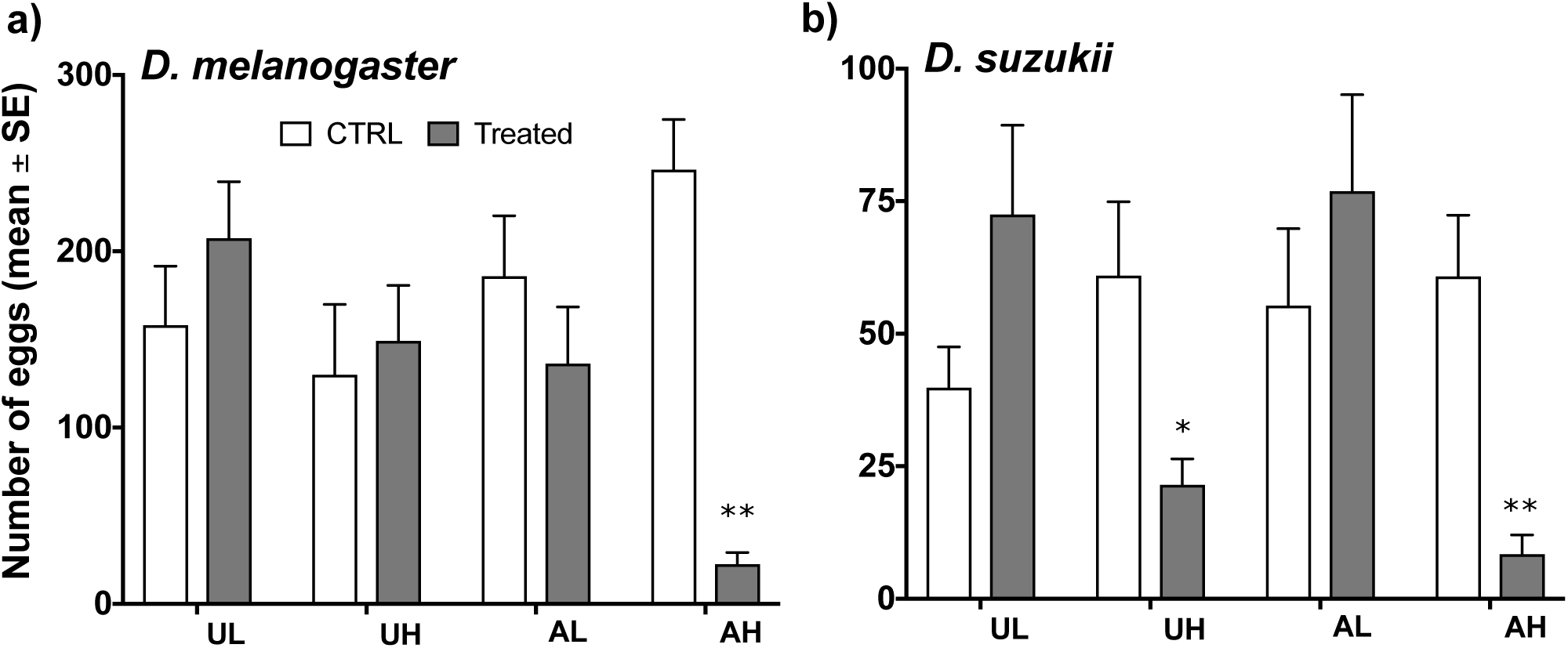
Oviposition site preference for standard food supplemented with dietary urea and ammonia (Treated) against non-supplemented standard food (CTRL). UL: standard food with 25 mM of urea; UH: standard food with 250 mM of urea; AL: standard food with 25 mM of ammonium chloride; AH: standard food medium with 250 mM of ammonium chloride. Mean ± standard error (SE) are shown. *=*p*<0.5, **=*p*<0.01.

**Figure 5.**
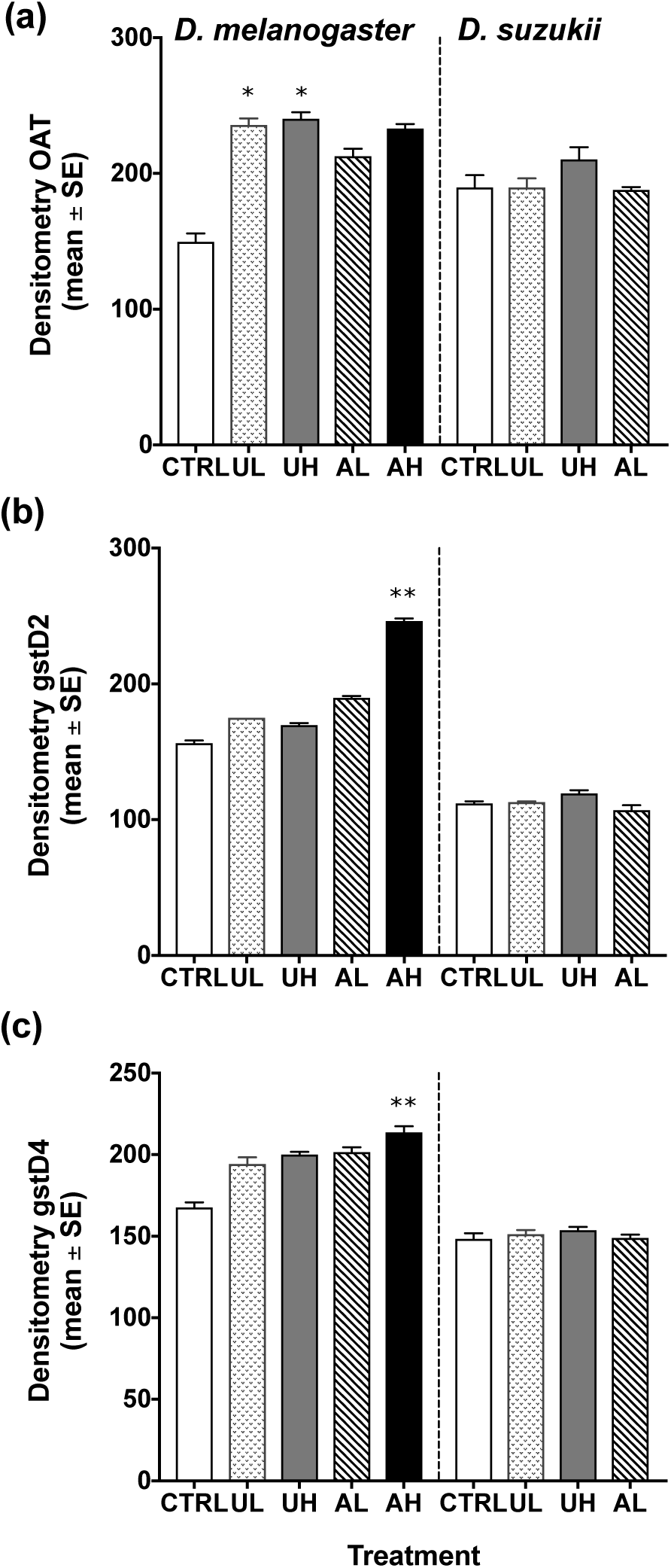
Semi-quantitative RT-PCR analysis of genes coding for OAT (a), gstD2 (b), and gstD4 (c) in *D. melanogaster* and *D. suzukii*, in control specimens (CTRL), in the presence of low (UL) and high (UH) concentrations of urea and in the presence of low (AL) and high (AH) concentrations of ammonium chloride. Data are mean ± standard error (SE). *=*p*<0.5, **=*p*<0.01.

### Expression of genes coding for metabolic and detoxifying enzymes

Semi-quantitative RT-PCR analysis of the expression of OAT, gstD2 and gstD4 evidenced a different expression pattern in the two species. Indeed, despite a common constitutive expression of the three genes in both the species observed in the control specimens, the expression of OAT, gstD2 and gstD4 resulted highly increased in *D. melanogaster* respect to *D. suzukii* after the exposure to ammonia and urea.

In particular, a significant increase of the OAT has been observed in *D. melanogaster* in the presence of urea, both at low and high concentration (*χ*^2^(4)=12.01, *p*<0.05), whereas no induction was evident in *D. suzukii* (*χ*^2^(3)=6.59, *p*=0.07; Figs. S3a, 5a). Differently from this pattern, gstD2 resulted highly induced in the presence of ammonium at high concentration in *D. melanogaster* (*χ*^2^(4)=13.06, *p*<0.001), whereas no significant difference has been detected at both concentrations of urea, neither for both treatments in *D. suzukii* (Figs. S3.b, 5b). Lastly, gstD4 showed an expression pattern similar to gstD2, with a significant induction under AH exposure in *D. melanogaster* (*χ*^2^(4)=13.08, *p*<0.001; Figs. S3.c, 5c).

## Discussion

We compared the effect of low and high concentrations of urea and ammonia on fecundity and larval development in *D. melanogaster* and *D. suzukii*. Our data show that fecundity is negatively affected in both species by nitrogenous waste products, but significantly greater effects have been observed in *D. suzukii*. When exposed to high concentrations of nitrogen compounds, female fecundity was more negatively affected by ammonia than by urea in both species, however, while *D. melanogaster* experienced a 50% fecundity reduction, *D. suzukii* fecundity was reduced by 70%.

Previous studies have documented the relevant role of ammonia as a sensory cue for female orientation and site selection in *D. melanogaster* (Delventhal *et al.* 2017; Min *et al.* 2013) and in other insect species (Bateman & Morton 1981; Kendra *et al.* 2005). Here, we have documented for the first time the greater sensitivity of *D. suzukii* pest to ammonia compared to *D. melanogaster*.

The stronger response observed in *D. suzukii* could derive from a greater olfactory and/or gustatory sensitivity to ammonia (e.g. Delventhal *et al.* 2017; Menuz *et al.* 2014). In fact, the concentration of volatiles associated with different maturation stages can greatly affect olfactory choices in *D. melanogaster* (e.g Versace *et al.* 2016; Zhu, Park & Baker 2003). Moreover, recent studies have shown how, during fruit maturation, changes occurring in the composition and concentration of volatiles can provide different cues for *D. suzukii* and *D. melanogaster* (Keesey, Knaden & Hansson 2015; Krause Pham & Ray 2015). Further studies should clarify the role of sensory cues in determining the greater sensitivity of *D. suzukii* to ammonia.

In addition, we observed a significant difference between species also regarding a typically non-volatile compound as urea, resulting in a greater reduction of fecundity in *D. suzukii* compared to *D. melanogaster*. While the number of laid eggs was only slightly decreased in *D. melanogaster* (Joshi *et al.* (1998) observed stronger effects at higher concentrations), *D. suzukii* experienced a 60% decrease in the number of laid eggs. This suggests again a higher repellent effect by nitrogenous waste in *D. suzukii*.

Interestingly, we observed differences in fecundity of *D. melanogaster* under urea exposure between choice (UH; 53% of total number of eggs) and no-choice (UH; 44% of total CTRL and UH eggs) assays suggesting that the presence of more cues or choices can ameliorate the inhibitory effect of urea on fecundity. This modulation though was not found in *D. suzukii* (choice, UH; 29% of total number of eggs; no choice, UH; 28% of total CTRL and UH eggs). The presence of a high level of ammonia instead caused a greater reduction in fecundity in the choice assay (*D. melanogaster*, 11%; *D. suzukii*, 9%) than in the no-choice assay (*D. melanogaster*, 37%; *D. suzukii*, 24%) in both species. Several studies have shown that *D. melanogaster* tends to hold eggs in absence of quality oviposition media (Joseph & Heberlein 2012; Schwarz, Durisko & Dukas 2014; Yang *et al.* 2008). However, *D. melanogaster* was found to lay eggs in substrate with potentially toxic chemicals when a harmless alternative is available (Azanchi, Kaun & Heberlein 2013).

We argue that the documented aversion of *D. suzukii* females for nitrogenous products might be an adaptive behavior to avoid substrates that can negatively affect larval fitness in this species more than in *D. melanogaster*. In fact, in *D. melanogaster* egg-laying behavior is influenced by the presence and density of larvae (Sarin & Dukas 2009; del Solar & Palomino 1966), and larval waste has been shown to modulate this effect (Aiken & Gibo 1979; Joshi *et al.* 1998). Along this line, we show that nitrogenous waste products affect *D. suzukii* larvae more negatively than *D. melanogaster* larvae. In our study, larval exposure to ammonia and urea resulted in high toxicity, showing a significant difference between species in the capacity to cope with the detrimental effects of these compounds. In *D. suzukii*, larvae were not able to survive in the presence of high concentration of ammonia, 100% of mortality was observed soon after hatching. On the other hand, larvae of *D. melanogaster* showed a delayed development, but viability remained comparable to the control. High levels of urea affected late larval stages in both species, interfering with larval survival as well as developmental time and pupation process, but with a stronger detrimental effect in *D. suzukii*.

Differently from many toxic chemicals that attack one or a few targets (Morton 1993; Russell *et al.* 1990), ammonia and urea are able to impact the whole organism (Cagnon & Braissant 2007; David *et al.* 1999; Yancey & Somero 1979). For this reason, we observe mechanisms to respond and resist the globally detrimental effects of these toxins. Strategies can involve uptake reduction, detoxifying pathways, as well as efficient excretory mechanisms (O’Donnell & Donini 2017). Interference with any of these processes could compromise an organism’ survival (Belloni & Scaraffia 2014; Scaraffia *et al.* 2005). Previous work showed that adaptation to urea and ammonia results in decreased larval feeding rate and longer developmental time in *D. melanogaster* (Borash *et al.* 2000; Botella *et al.* 1985). This could explain our data at high levels of ammonia, where a significant increase in developmental time was associated with larval viability in *D. melanogaster*. However, in *D. suzukii* compensation mechanisms failed, causing ammonia levels to rapidly increase, resulting in an acute intoxication. This hypothesis is supported by significant metabolic and detoxification differences observed between the two species, where ornithine aminotransferase as well as glutathione-S-transferase gene expression increased under high levels of ammonia and urea in *D. melanogaster*, whereas no induction was evident in *D. suzukii*. The ornithine aminotransferase enzyme plays a crucial role in amino acids metabolism and nitrogen homeostasis (Ventura *et al.* 2008; Yoshida, Juni & Hori 1997), while glutathione-S-transferase is highly expressed in response to xenobiotics and oxidative stress (Hamby *et al.* 2013; Nguyen *et al.* 2016). Alteration in their expression can be associated with inefficient detoxification and reduction of tolerance capacity to environmental stressors (Claudianos *et al.* 2006; Nguyen *et al.* 2016; Passador-Gurgel *et al.* 2007; Seiler 2000). This explains the mortality observed in *D. suzukii* when first instar larvae were faced with high level of ammonia in the medium. Urea can interfere with important cell processes, act as protein denaturant, and reduce enzyme activity (Somero & Yancey 1997; Yancey 1992; Yancey & Somero 1979), resulting in development delay and larval stop (Botella *et al.* 1985; Shiotsugu *et al.* 1997). Drosophila species are not familiar with this compound, due to the fact that they are not able to produce it, suggesting a lack of physiological mechanism to handle it. However, we observed a similar enzymatic response under urea and ammonia exposure, in agreement with previous studies describing the evolution of cross-tolerance between these stress traits (Borash *et al.* 2000; Borash & Shimada 2001). Developmental delay and reduction in feeding rate are adaptive strategies developed to reduce urea uptake in *D. melanogaster* larvae and favour toxin resistance (Borash *et al.* 2000; Botella *et al.* 1985; Etienne, Fortunat & Pierce 2001). Larvae of *D. suzukii* are characterized by a longer developmental time to reach the maximum size and to enter the pupal stage (Hamby *et al.* 2016; Wegman, Ainsley & Johnson 2010). A delay combined with urea alteration of protein synthesis (Somero & Yancey 1997) could strongly compromise eggs-to-pupae viability. This effect combined with a lack of efficient detoxifying mechanisms confirmed by our study, explains well the incapacity *D. suzukii* to cope with high load of urea.

In the wild, the adaptation of *D. suzukii* to fresh fruits as oviposition substrate, has allowed larvae to develop in a safer and healthier environment. However, our study shows how metabolic adaptations to fresh food have resulted in less efficient detoxifying and excretory mechanisms. Further studies are necessary to better understand the interaction between female fecundity and nitrogenous compounds for possible use of these chemicals as repellants or reproductive toxicants in natural conditions. Moreover, more attention needs to be directed to the larval stage in *D. suzukii*, which is highly affected by environmental variations and shows important adaptations to a diverse ecological niche with respect to *D. melanogaster*. Our findings, in fact, suggest the possibility to significantly compromise larval fitness and survival in the pest species via the exposure to environmental compounds.

## Authors’ Contributions

VB conceived the study and designed methodology; VB, AG, GB and MM collected the data; VB and AG and MM analysed the data; VB drafted the manuscript; VB, EV, AH and MM led the writing of the manuscript; EV and AH supervised the study. All authors contributed critically to the drafts and gave final approval for publication.

## Acknowledgements

VB was funded by the grant “Sustainable strategies against pest fruit flies: from olfactory perception to the protection of the small fruits of Trentino” of the City of Rovereto, EV was supported by a Harvard Mind Brain and Behavior Faculty Award to Benjamin de Bivort.

